# Bilateral Asymmetry of the Forearm Bones as Possible Evidence of Antemortem Trauma in the StW 573 *Australopithecus* Skeleton from Sterkfontein Member 2 (South Africa)

**DOI:** 10.1101/486076

**Authors:** A.J. Heile, Travis Rayne Pickering, Jason L. Heaton, R.J. Clarke

## Abstract

The 3.67-million-year-old StW 573 *Australopithecus* skeleton is important for the light it sheds on the paleobiology of South African species of that genus, including, as discussed here, how the possible pathology of the specimen informs our understanding of *Australopithecus* behavior. The StW 573 antebrachium exhibits bilateral asymmetry, with significantly more longitudinally curved left forearm bones than right. Arguing from a comparative perspective, we hypothesize that these curvatures resulted from a fall onto a hyperextended, outstretched hand. It is unlikely that the fall was from a significant height and might have occurred when the StW 573 individual was a juvenile. This type of plastic deformation of the forearm bones is well-documented in modern human clinical studies, especially among children between the ages of four and ten years who tumble from bicycles or suffer other common, relatively low-impact accidents. Left untreated, such injuries impinge normal supination and pronation of the hand, a condition that could have had significant behavioral impact on the StW 573 individual.

## Introduction

The 3.67-million-year-old StW 573 (“Little Foot”) *Australopithecus* skeleton, from Member 2 of the Sterkfontein Formation (South Africa), was discovered by R.J. Clarke in a sequence of steps that unfolded in the laboratory and field over the years of 1995 through 1997 (see, e.g., Dugard, 1995; Clarke, 2018). In a series of papers, Clarke and his colleagues (Clarke and Tobias, 1995; Clarke, 1998, 1999, 2002, 2007, 2008, 2018; Pickering et al., 2004; Granger et al., 2015; Stratford et al., 2017; Beaudet et al., 2018a,b; Clarke, 2018; Clarke et al., 2018; Crompton et al., 2018; Heaton et al., 2018) have provided information on the skeleton’s stratigraphic and taphonomic circumstances, geochronological position, taxonomic status, and functional morphology.

The last set of analyses conclude that the “Little Foot” individual (from here, LF), when alive, was an orthograde terrestrial biped and also possessed significant adaptations for climbing (e.g., Clarke and Tobias, 1995; Beaudet et al., 2018b; Carlson et al., 2018; Crompton et al., 2018; Heaton et al., 2018). It is in this context that the observed asymmetry of the LF skeleton’s right (lacking significant diaphyseal curvature) and left (displaying significant diaphyseal curvature) forearm bones (Heaton et al., 2018) is of particular interest (Fig 1).^1^

**Figure 1.**
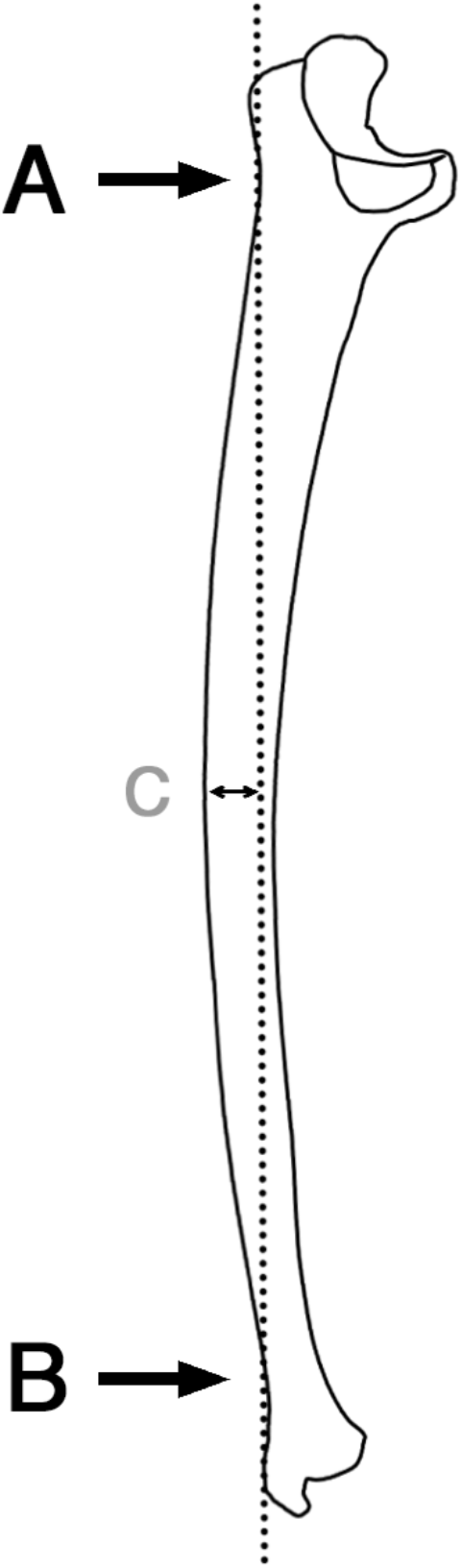
Sketch of a *Pan troglodytes* right ulna shown in lateral view to illustrate calculation of curvature subtense, where A indicates the inflection point on the posterior margin of the ulna at the level of the radial notch, B indicates a second inflection point on the posterior margin of the ulna at the minimal distal circumference, and C indicates the maximum distance from the posterior-most margin of the ulna to the line drawn between A and B (Aiello et al. 1999; Drapeau et al. 2005). Figure modified from Drapeau et al. (2005: 618, Fig. 16).

Like other diaphyseal bone curvatures in primates (e.g., Susman, 1979; Bruns et al., 2002; Richmond, 2007), those of the radius (lateral) and ulna (dorsal) are probably the result of both genetics and limb function. As to the latter, biomechanical studies suggest that radial lateral bending is, in part, a plastic response to the exertion of the pronator teres during pronation (Galtés et al., 2009), while it is widely hypothesized that ulnar dorsal curvature results from the powerful engagement of both pronator and supinator muscles and as an adaptation that expands the interosseous membrane for insertion of enlarged digital flexors (e.g., Miller, 1933; Knussmann, 1967; Swartz, 1990). In this hypothetical context, it is notable that below-branch suspensory locomotorists show significantly curved radii and ulnae (e.g., Knussmann, 1967).

Based on muscle weight, bone weight, and osteometrics, forelimb bone asymmetries have been documented in several extant primates (e.g., Schultz, 1937; Dhall and Singh, 1977; Falk et al., 1988; McGrew and Marchant, 1997). Further, in a study of the total subperiosteal areas of wild-caught chimpanzee (*Pan troglodytes*) humerus diaphyses, Hunt’s (1991) observations on that animal’s positional behavior was cited to suggest that bilateral asymmetry of this element might be due to unimanual suspension (Sarringhaus et al., 2005). Thus, it is not unreasonable to predict a similar bilateral asymmetry of forelimb bone curvatures in hominoids that regularly engage in below-branch positions and activities. To our knowledge, no published data are available to test this prediction. However, anecdotally, we have not observed significant bilateral curvature asymmetries in the antebrachial segments of chimpanzee and bonobo (*Pan paniscus*) skeletons that we have studied.

It seems as likely, if not more, that the differential morphology of the right and left forearm bones of the LF skeleton are due to injury than to normal *in vivo* functional demands of its forelimbs. Specifically, we hypothesize that, as a juvenile, the LF individual suffered acute plastic bowing deformation of its left forearm. We base this hypothesis on the greater likelihood that it is the curved, left, and not the straight right, antebrachial elements of the LF individual that are abnormal. We argue this point from both observational and comparative perspectives. First, there is no apparent antemortem trauma to the right humerus, radius, or ulna of LF (Heaton et al., 2018). In addition, the bones display no sign of atrophy, which is predicted if the right upper limb was in disuse during the life of LF. Second, survey of the literature on forearm bone curvature in non-*Homo* late Pliocene and early Pleistocene hominins predicts that a small-bodied *Australopithecus*, such as LF, should possess relatively straight radii and ulnae.

If our hypothesis is correct, then the curved left antebrachial elements of the LF skeleton stand as some of the earliest evidence of traumatic antemortem skeletal injury in the hominin^2^ fossil record. Plastic deformation of the arm requires longitudinal forces within a limited range of magnitude. These forces exceed those causing temporary elastic deformation and approach fracture-inducing forces; the resulting bowing remains unless physically corrected (Borden 1974, 1975). It is hypothesized here that the dorsal diaphyseal curvature evidenced in LF’s left forearm represents traumatic bowing from a fall onto a hyperextended, outstretched hand. This likely occurred when LF was young, as mature bone is more prone to fracture catastrophically, rather than bow (Borden 1975).

## Materials and Methods

The goal of the first part of our analysis of the antebrachial asymmetry of LF is to place its radial and ulnar curvatures in comparative fossil context. A widely used method for representing forearm bone curvature in paleoanthropology is the calculation of curvature subtenses (e.g., Aiello et al. 1999) (Fig. 2). No matter the measurement(s) of curvatures, though, as discussed above, it seems that they can be induced by a number of different processes or combination of processes. Among those causes is allometry, as it is generally appreciated that the more robust an ulna is anteroposteriorly, the greater its curvature will be (e.g., Drapeau et al., 2005).

**Figure 2.**
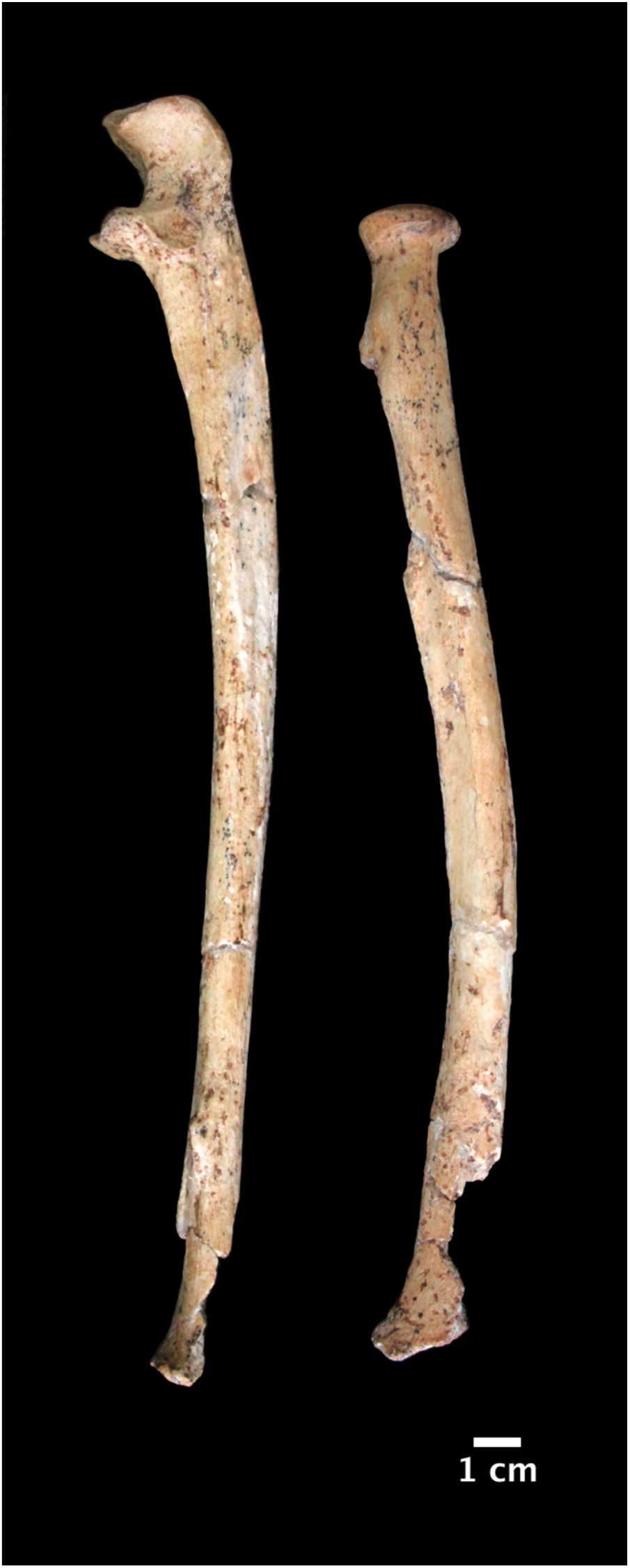
The curved left forearms bones of the StW 573 *Australopithecus* skeleton shown with superior toward the top of the image. The ulna (left) is near-lateral view and radius (right) is in anterior view.

With this understanding, we consulted the literature to obtain length, anteroposterior midshaft diameter, and curvature subtense values (terms defined in Table 1) for relevant late Pliocene and early Pleistocene non-Homo hominin ulnae (Drapeau et al., 2005; Churchill et al., 2013) in order to compare those values to those of the left ulna of LF (Heaton et al., 2018). Relevant exceptions to this were the anteroposterior midshaft diameter and curvature subtense of the *Australopithecus sediba* MH2 ulna, which we calculated from published photographs (Rein et al., 2017) using image analysis software. We omitted radius measurements from our study because of the paucity of complete and near-complete *Australopithecus* and *Paranthropus* comparative radii. Next, in order to contextualize these fossil results, we reviewed the clinical literature on traumatic forearm bone bowing of modern humans.

**Table 1.**
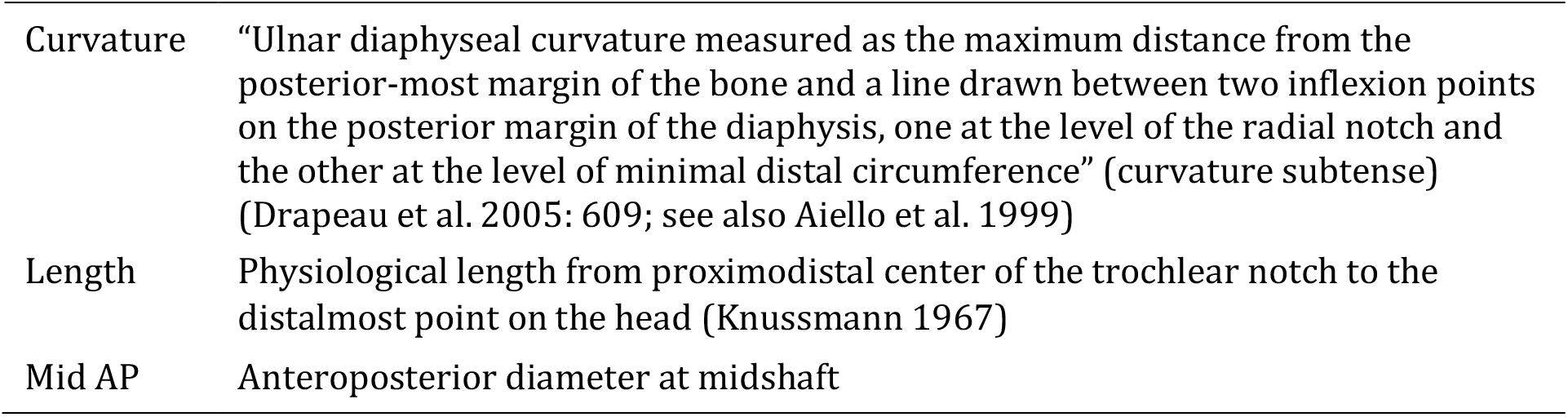
Ulna measurement abbreviations and definitions used in this study

|  |  |
| --- | --- |
| Curvature | “Ulnar diaphyseal curvature measured as the maximum distance from the posterior-most margin of the bone and a line drawn between two inflexion points on the posterior margin of the diaphysis, one at the level of the radial notch and the other at the level of minimal distal circumference” (curvature subtense) (Drapeau et al. 2005: 609; see also Aiello et al. 1999) |
| Length | Physiological length from proximodistal center of the trochlear notch to the distalmost point on the head (Knussmann 1967) |
| Mid AP | Anteroposterior diameter at midshaft |

**Table 2.** Measurements of complete and near-complete *Australopithecus* and *Paranthropus* ulnae^a b^

| Genus | Specimen | Length<sup>c</sup> | Mid AP<sup>d</sup> | Curvature<sup>e</sup> |
| --- | --- | --- | --- | --- |
| *Australopithecus* | AL 288-1 | 181/206 | 12.3 | 2.5/4.1 |
|  | AL 438-1 | 252 | 17.6 | 7.5 |
|  | MH2 | 233 | [11.7] | [4] |
|  | StW 573 | 244 | 14.2 | [12] |
| *Paranthropus* | L40-19 | 295 | 14.8 | 12.5 |
|  | OH 36 | 247/277 | 17.3 | 13.9/15.7 |

<sup>a</sup>Measurements described in Table 1. All measurements in millimeters. Values in brackets represent estimates
<sup>b</sup>For taxonomic assignments, see Johanson et al. (1982), Howell et al. (1987), Aiello et al. (1999), Clarke (1998), Drapeau et al. (2005), and Berger et al. (2010)
<sup>c</sup>Data for AL 288-1, AL 438-1, L40-19, and OH 36 from Drapeau et al. (2005); MH2 from Churchill et al. (2013); StW 573 from Heaton et al. (in submission)
<sup>d</sup>Data for AL 288-1, AL 438-1, L40-19, and OH 36 from Drapeau et al. (2005); MH2 estimated from image software using scaled photo from Rein et al. (2017); StW 573 from Heaton et al. (in submission)
<sup>e</sup>Data for AL 288-1, AL 438-1, L40-19, OH 36 from Drapeau et al. (2005); MH2 estimated from image software using scaled photo from Rein et al. (2017); StW 573 from Heaton et al. (in submission)

## Results and Discussion

The left ulna of LF displays a comparatively high estimated curvature value of 12, much higher than those of other measurable *Australopithecus* ulnae and nearer to the curvature values of the early Pleistocene hominin ulnae OH 36 (Olduvai Gorge, Tanzania) and L40-19 (Omo, Ethiopia) (Table 2). Like OH 36 and L40-19, the left ulna of LF is relatively large. The large size of these elements may account, at least in part, for their significant curvatures (e.g., Knussmann, 1967; Drapeau et al., 2005). In this context, it is not irrelevant to note that the diaphysis of the largest known radius in the early hominin fossil record, OH 80, from Olduvai, is also apparently strongly curved (Domínguez-Rodrigo et al., 2013).

However, it is also possible that the extreme curvatures of OH 36, L40-19, and OH 80 are taxonomic-specific, caused by genetics and/or locomotor adaptations (such as forelimb-dominated climbing). All of these specimens have been assigned (at least tentatively) to the genus *Paranthropus* (e.g., Howell and Wood, 1987; Aiello et al., 1999; Domínguez-Rodrigo et al., 2013). If that is the case, then the curved ulna of LF is particularly intriguing given that it derives from an *Australopithecus* individual,^3^ and that even large *Australopithecus* ulnae, such as AL 438-1, are only moderately curved (Table 2). And, although there are fewer complete or near-complete *Australopithecus* radii than ulnae, they, too, seem to show rather straight diaphyses, more comparable to diaphyseal profiles of modern human forearm bones (e.g., Johanson et al., 1982; Heinrich et al., 1993; Toussaint et al., 2003; Pickering et al., 2018).

In sum, the curvature of the left antebrachium of LF is unexpected in a comparative fossil context and seems to suggest that it is this side that is abnormal and not the straight, right forearm bones of the skeleton. As discussed in the introduction, we have been unable to find data on modern primates to suggest that asymmetrical bowing of the forearm bones results from unimanual locomotor, positional, or other behaviors. Thus, it is most parsimonious to hypothesize that the curvature of the left forearm bones of LF resulted from an antemortem trauma.

There is a large body of experimental work on plastic deformation of mammalian long limb bone diaphyses (e.g., Chamay 1970; Chamay and Tschantz 1972; Lanyon, 1980; Lieberman et al. 2003). Traumatic bowing fracture (TBF) is a specific type of plastic deformation that, in modern human clinical circumstances, most often affects the bones of the antebrachium. It presents as a distinct abnormal curvature of the limb bone diaphysis, with microfracturing on the concave side of the bow (e.g., Borden 1974, 1975). Though rare, adult cases of TBF of the forearm bones have been documented and usually result from slow bending forces such as those caused by entrapment in rotating machines (e.g., Greene 1982; Simonian and Hanel 1996; Sclamberg et al. 1998; Sen et al., 2004; Lefaivre et al. 2007; Tada et al., 2008; Tianhao et al., 2014). However, most often, it is children between the ages of four and ten years that experience TBF (Vorlat and De Boeck 2003) as a result of falls onto outstretched, hyperextended hands (e.g., Borden 1974, 1975; Naga and Broadrick 1977; Crowe and Swischuk 1977; Aponte and Ghiatas 1989; Vorlat and De Boeck 2003; Musters and Colaris 2017). Descriptions of TBF in children vary in their level of detail, but it appears as though most falls that result in bowing are from rather insignificant heights and are, at least sometimes, associated with varying degrees of momentum. For instance, bicycle accidents are a commonly reported cause of injury (e.g., Borden 1974, 1975; Naga and Broadrick 1977; Crowe and Swischuk 1977; Komara et al. 1986).

## Conclusion

The fossil record of late Pliocene and early Pleistocene African hominins includes several excellently preserved partial skeletons, including the *Australopithecus* specimens A.L. 288-1 (“Lucy,” Hadar, Ethioipia), KSD-VP-1 (“Kadanuumuu,” Woranso-Mille, Ethiopia), DIK-1-1 (“Selam,” Dikika, Ethiopia), Sts 14 and StW 431 (Sterkfontein), and MH1 and MH2 (Malapa, South Africa), as well as the *H. ergaster*“Nariokotome Boy” (KNM-WT 15000) from West Turkana, Kenya. Research on these, and on other less complete specimens, has revealed various lines of evidence related to the peri- and postmortem fate of our early ancestors. Those data support hypotheses that early hominins, like modern humans, suffered various kinds of fatal misfortune, including succumbing to the predation of carnivores (e.g., Brain, 1981, Pickering et al., 2004), raptors (Berger and Clarke 1995), and possibly crocodilians (Njau and Blumenschine, 2012; contra, Baquedano et al., 2012), to falls from significant heights (L’Abbé et al., 2015; Kappelman et al., 2012), and to septicemia (Walker, 1993).

And, although there have been several interesting observations of early hominin dental and periodontal defects (e.g., Robinson, 1956; White, 1978; Tobias, 1991; Walker, 1993; Ripamonti et al., 1997; Guatelli-Steinberg, 2003), abnormal bone apposition (Walker et al., 1982; Rothschild et al., 1995), and bony pathologies (Cook et al., 1983; Latimer and Ohman, 2001; D’Anastasio et al., 2009; Domínguez-Rodrigo et al., 2012), less is known about traumatic antemortem damage to skeletons of this great antiquity. It has been proposed that the ~3.6 Ma Kadanuumuu skeleton shows evidence of an unhealed fibular fracture, incurred when the individual was a juvenile (Lovejoy et al., 2016), and the ~3.0 Ma *Australopithecus afarensis* A.L. 333-107 humerus, from Hadar might preserve a healed proximal fracture (Lovejoy et al., 1982). With the likely acute plastic bowing deformation of its left antebrachium, the LF skeleton joins the last group as a possible example of skeletal trauma incurred during the life of an important early hominin specimen.

If indeed the curvature exhibited in LF’s left antebrachium represents a TBF, this has possible implications for the ways in which the individual interacted with the environment. First, if modern humans serve as a reasonable referent in this case, then it is more likely than not that LF’s injury occurred prior to adulthood. As discussed above, most descriptions of the causes of childhood TBF are vague, but, based on that literature, it seems most reasonable to suggest that the injury was incurred under low-impact conditions. In this sense, the severity of the suggested LF injury is unlike the trauma proposed to have been incurred by Lucy. Kappelman et al. (2016) report on possible greenstick fractures of multiple skeletal elements of Lucy and suggest these injuries resulted from a fall of great heights. They propose these fractures likely caused damage to Lucy’s internal organs, resulting in her death, and hypothesize the fall was from a tree, thus offering evidence for arborealism in *A. afarensis*. Although some of the LF morphology is reflective of arboreal locomotor behavior (Clarke and Tobias 1995; Beaudet et al., 2018b; Crompton et al., 2018), the proposed LF injury cannot be used in support of arborealism, as TBF can occur at ground level.

In the case of modern humans, TBFs can result in the loss of full pronation and supination unless corrected (e.g., Borden 1974, 1975; Greene 1982; Sanders and Heckman 1984; Anderson et al., 1994). It seems safe to assume that LF did not receive corrective treatment for TBF. That said, permanent unimanual limitation of forearm rotation may have hindered her climbing and manipulatory abilities, which is significant for an organism that otherwise appears to have been reasonably adapted to climbing (Clarke and Tobias 1995; Beaudet et al., 2018b; Crompton et al., 2018) and that possessed a hand that shows at least some modern humanlike characteristics (Clarke 1999). That the LF individual lived to an advanced age (Clarke et al., 2018) with such a possible handicap raises obvious questions about sociality and intraspecific care in South African *Australopithecus*.

## Acknowledgements

This work was funded by a Kellet Mid-Career Award from the University of Wisconsin-Madison (USA) awarded to TRP and by additional funds from the National Research Foundation (South Africa) awarded separately to Dominic Stratford and Kathleen Kuman. Thanks to Abel Molepolle and Andrew Phaswana for their continued assistance and camaraderie, as well as the rest of the Little Foot research team, Amelie Beaudet, Laurent Bruxelles, Kristian Carlson, Robin Crompton, Tea Jashashvili, Kathleen Kuman, Juliet McClymont, and Dominic Stratford.

## Footnotes

1 The differential maximum lengths of the LF radii (left = 236 mm; right estimate = 250 mm; Heaton et al., 2018) also reflects this variance in curvature.

2 Based on White (2002), Clarke (2012), and White et al. (2015), RJC objects to the use of the term “hominin,” and prefers the use of the term “hominid.”

3 Taxonomic determination of LF is based primarily on its craniodental anatomy (Clarke et al., 2018). However, we also note that that the left ulna of the specimen also shows typical *Australopithecus* features (Heaton et al., 2018), such as an anteriorly directed and minimally keeled trochlear notch (Aiello et al., 1999; Drapeau 2004, 2008).

